# Batch Normalization Followed by Merging Is Powerful for Phenotype Prediction Integrating Multiple Heterogeneous Studies

**DOI:** 10.1101/2022.09.28.509843

**Authors:** Yilin Gao, Fengzhu Sun

## Abstract

Heterogeneity in different genomic studies compromises the performance of machine learning models in cross-study phenotype predictions. Overcoming heterogeneity when incorporating different studies in terms of phenotype prediction is a challenging and critical step for developing machine learning algorithms with reproducible prediction performance on independent datasets. We investigated the best approaches to integrate different studies of the same type of omics data under a variety of different heterogeneities. We developed a comprehensive workflow to simulate a variety of different types of heterogeneity and evaluate the performances of different integration methods together with batch normalization by using ComBat. We also demonstrated the results through realistic applications on six colorectal cancer (CRC) metagenomic studies and six tuberculosis (TB) gene expression studies, respectively. We showed that heterogeneity in different genomic studies can markedly negatively impact the machine learning classifier’s reproducibility. ComBat normalization improved the prediction performance of machine learning classifier when heterogeneous populations presented, and could successfully remove batch effects within the same population. We also showed that the machine learning classifier’s prediction accuracy can be markedly decreased as the underlying disease model became more different in training and test populations. Comparing different merging and integration methods, we found that merging and integration methods can outperform each other in different scenarios. In the realistic applications, we observed that the prediction accuracy improved when applying ComBat normalization with merging or integration methods in both CRC and TB studies. We illustrated that batch normalization is essential for mitigating both population differences of different studies and batch effects. We also showed that both merging strategy and integration methods can achieve good performances when combined with batch normalization. In addition, we explored the potential of boosting phenotype prediction performance by rank aggregation methods and showed that rank aggregation methods had similar performance as other ensemble learning approaches.

**Author summary:** Overcoming heterogeneity when incorporating different studies in terms of phenotype prediction is a challenging and critical step for developing machine learning algorithms with reproducible prediction performance on independent datasets. We developed a comprehensive workflow to simulate a variety of different types of heterogeneity and evaluate the performances of different integration methods together with batch normalization by using ComBat. We also demonstrated the results through realistic applications on six colorectal cancer (CRC) metagenomic studies and six tuberculosis (TB) gene expression studies, respectively. From both the simulation studies and realistic applications, we showed that batch normalization is essential for improving phenotype prediction performance by machine learning classifiers when incorporating multiple heterogeneous datasets. Combined with batch normalization, merging strategy and ensemble weighted learning methods both can boost machine learning classifier’s performance in phenotype predictions. In addition, we explored that rank aggregation methods should be considered as alternative way to boost prediction performances, given that these methods showed similar robustness as ensemble weighted learning methods.

## Introduction

Genotype to phenotype mapping is an essential problem in the current genomic era. With the development of advanced biotechnologies, many types of genomic data such as single nucleotide polymorphisms, gene expression profiles, proteomics, metagenomics, etc. have been generated in many different studies. These omics data provide essential resources to understand the relationships between omics data and phenotypes. Despite these fundamental developments, due to the heterogeneity of data, it is challenging to integrate the omics data to understand genotype to phenotype mapping. For a single type of data such as gene expression or metagenomic data, many sources of heterogeneity can occur. For example, the samples can come from different ethnic groups with varying underlying distributions of the features. Even if the samples come from the same population, the genomic data can be generated from different laboratories and/or derived from different experimental technologies resulting in different distributions of the data. Another types of heterogeneity can be caused by the different causal mechanisms of the same phenotype in the populations under study [1]. The objective of this study is to investigate the best approaches to integrate different studies of the same type of data under a variety of different heterogeneities. In this work, we concentrate on gene expression profiles or microbial abundance in metagenomic studies.

Many machine learning algorithms including linear regression, logistic regression, penalized regression, support vector machines (SVM), random forests (RF), neural networks (NN) and deep neural networks (DNN) have been used to predict phenotypes from omics data [2–5]. Most previous studies validated the prediction methods using within dataset cross validation usually with relatively high prediction accuracy.

However, the prediction accuracy is markedly decreased when the learned algorithms are used in independent datasets [6, 7]. Many sources of study heterogeneity, for example, different experimental platforms or procedures and differences in patient cohorts [1], all contribute to compromise the prediction performance of machine learning models in cross-study settings. Thus, overcoming heterogeneity in cross-study phenotype prediction is a critical step for developing machine learning algorithms with reproducible prediction performance on independent datasets.

Many studies have been carried out to mitigate the heterogeneity in cross-study phenotype predictions. Zhang et al. [8] focused on the batch effects of data when developing genomic classifiers. Patil et al. [9] simulated genomic samples and perturbed the coefficients of linear relations between outcomes and predictors to evaluate model reproducibility with different degrees of heterogeneity. In this study, we address three types of heterogeneity: different background distributions of genomic features in populations, batch effects across different studies from the same population, and different disease models in various studies. We aim to evaluate how different statistical methods can mitigate these three types of heterogeneity.

Merging all datasets into one and treating all samples as if they are from the same study is a generally used method for cross-study predictions. With the increase of sample size and diversity in the study population, merging method has been shown to lead to better prediction performance than using only individual studies [2, 5, 10]. Another approach is to integrate the trained predictors from different machine learning models derived from various training datasets. Ensemble weighted learning is a commonly used integration method to deal with the impact of heterogeneity on cross-study prediction performance. Ensemble learning methods that integrate predictions from multiple machine learning models showed the ability to boost the prediction performance than using only the component methods that the ensemble learning contains [9, 11]. Besides ensemble weighted learning methods, aggregating the ranks from sample predicted probability instead of the probability itself offers a promising alternative for integration. In some situations, the predicted probabilities for the samples in the test data may not be correct, but the relative order could provide some useful information. In such situations, aggregating the ranks instead of the predicted probabilities might be more reasonable. To the best of our knowledge, no studies investigated rank aggregation methods based on omics data phenotype prediction.

ComBat [12] normalization is a commonly used method for removing batch effects between different datasets. In our previous study [13], we showed that when dealing with heterogeneity in cross-study predictions, applying ComBat only before training machine learning models did not improve the prediction performance. Zhang et al. [8] showed that ensemble weighted learning methods outperform batch correction by ComBat at high level of batch differences. Nevertheless, in this study, we aim to explore the potential of combining the normalization of ComBat together with merging and integration methods (ensemble weighted learning and rank aggregation) in the presence of three different types of heterogeneity mentioned above. We provide both simulations and real data applications on metagenomic and gene expression data to show the comparisons of performance from different statistical methods when dealing with cross-study heterogeneity.

## Methods

### Outline of workflow for integrating multiple heterogeneous metagenomic datasets

To investigate the prediction performance of different merging and integration methods when applied to multiple heterogeneous datasets, we developed a comprehensive workflow with three main steps to conduct the experiments.

The first step of our workflow is to simulate heterogeneous metagenomic datasets under three different scenarios as shown in Fig 1A. In the first scenario, we investigated the impacts of different background distributions of genomic features in populations on the machine learning model’s prediction performance. Due to population differences such as different ethnicity, diet, etc, the background distributions of the genomic features such as SNPs, expression levels and microbial abundance in microbiome can be different. Thus, when training machine learning classifiers on one population and predict on a different population, the heterogeneity in the background distributions may negatively impact the prediction performance if the heterogeneity between the training and test datasets is ignored. In order to determine how the population difference heterogeneity can affect machine learning classifier’s performance as well as find the best approaches for integrating different prediction methods from several heterogeneous studies, we simulated three different populations with different background genomic distributions and manipulated the differences. The detailed implementation of simulating the two training datasets and one test dataset is described in Scenario 1: Different background distributions of genomic features in populations.

**Fig 1.**
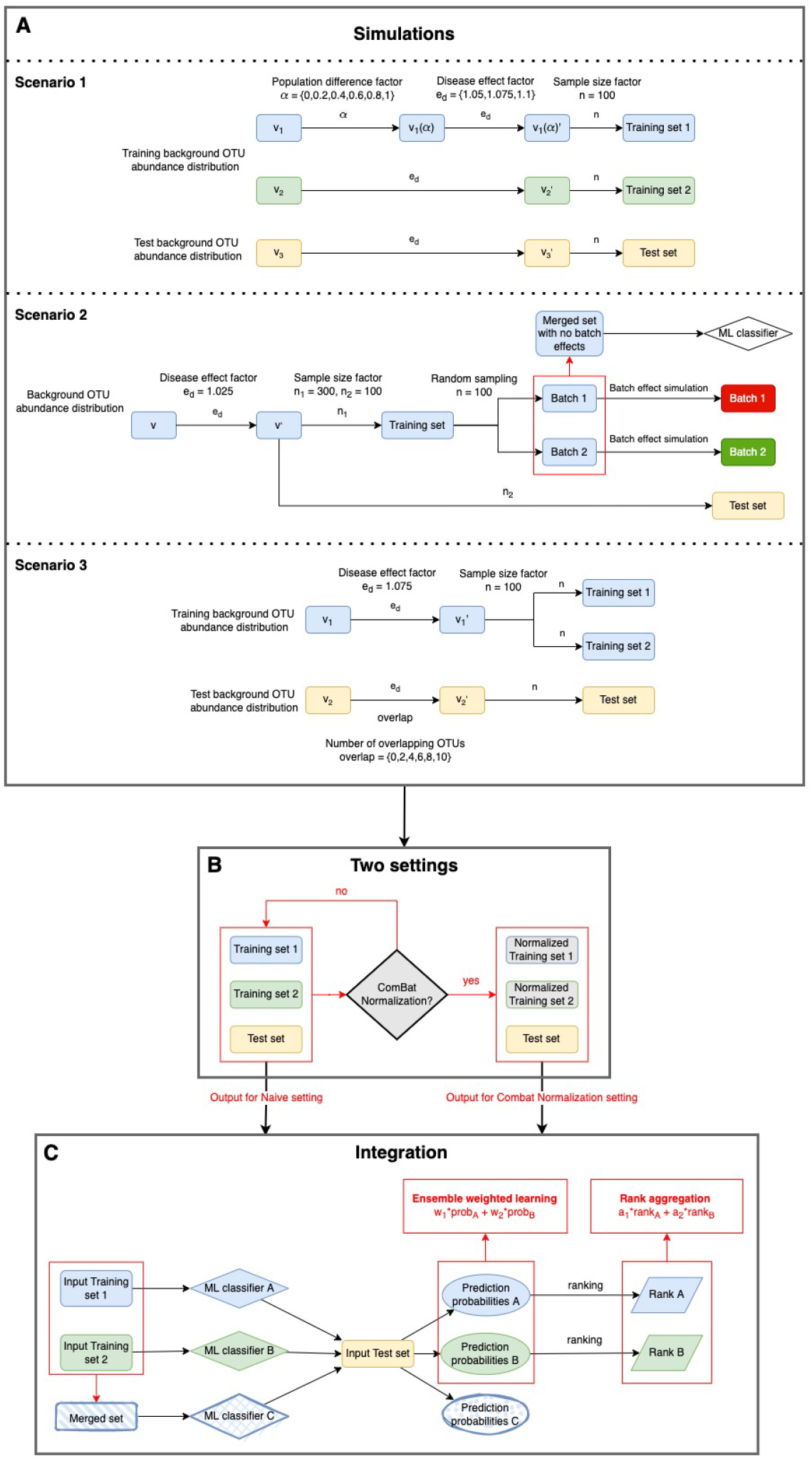
Workflow for integrating multiple simulated heterogeneous metagenomic datasets. A: Simulation step of three different heterogeneity scenarios. Scenario 1: Different background distributions of genomic features in populations. Scenario 2: Different batch various studies. The output from this step consists of two simulated training datasets and one test dataset. B: Naive and ComBat normalization settings in this study. The output from A are directly used in naive setting, while the two training datasets in the output are normalized first in ComBat normalization setting. C: Integration step of applying ensemble weighted learning and rank aggregation methods. The output training datasets from the previous step are used in training machine learning classifiers. Then the predictors are integrated by ensemble weighted learning methods or the ranks of the predictors are integrated by rank aggregation methods.

The second scenario of heterogeneity is related to batch effects. Batch effects are referred to as non-biological variations across different batches of data, typically, variations caused by technical differences across experimental conditions or laboratories. It has been argued that the reproducibility of genomic findings can be weakened by batch effects [14]. Batch effect correction is a common pre-processing step when dealing with genomic data. Many batch effect correction methods have been proposed recently, such as ComBat [12], edgeR [15] and DESeq2 [16]. Several studies proposed ensemble learning methods that can potentially mitigate batch effects [8, 9]. Here we investigated how those methods perform in terms of mitigating batch effects on prediction performance of binary classifiers through simulation studies. In this scenario, the training and test datasets are from the same background distribution of genomic features. However, after the simulation of training and test datasets, batch effects were simulated on training datasets to form two different batches. These two batches were served as the training datasets in the following experiments. The detailed method is described in Scenario 2: Different batch effects in studies with the same background distribution of genomic features in a population.

In the above two scenarios, we considered the effects of different background distribution of genomic features in populations and batch effects on the classifier’s performance, and we assumed that the underlying disease models for either the same or different populations are the same. However, several studies have shown that the associated microbes can potentially be population dependent. For example, CRC development varies in populations with different status of diabetes [17], smoking [18] and obesity [19]. Thus, we further explored the performance of those merging and integration methods when the underlying disease models are different. More specifically, the disease related gnomic features are different in training and test datasets. This is the third scenario in our workflow. We tuned the number of overlapping disease related microbes in training and test disease model, and simulated different datasets accordingly. The description of methods can be found in Scenario 3: Different disease models in different studies

After the simulations of training and test datasets from the above three scenarios as shown in Fig 1A, we aimed to evaluated the performance of merging and integration methods in two settings as shown in Fig 1B: naive setting with direct use of the simulated training and test datasets from the first step into the third integration step, and ComBat normalization setting with normalizing the two training datasets before applying the integration and merging methods. In the Methods section Naive and ComBat normalization settings, we described how the ComBat normalization was conducted on the training datasets in details.

Then, two simulated training datasets and one test dataset from the above steps are served as input of the last step in this workflow. In this step, a machine learning classifier was trained on each training dataset independently, and then different integration methods were applied on the two predictors and generated final results. The predictors from the trained machine learning classifiers were directly used in ensemble weighted learning methods, while when applying rank aggregation methods, the predicted probabilities were converted to ranks. A more detailed description of the ensemble weighted learning and rank aggregation methods we used is in Methods. In addition to these integration methods, we also applied the merging strategy. As shown in Fig 1C, we pooled the two training datasets together and trained machine learning classifier on the merged dataset, then predicted on the test dataset directly to get final results. The performances of merging and integration methods were compared later. In this study, we chose random forests (RF) [20] as the machine learning classifier to be used in the above procedures.

### Simulation strategies

#### Scenario 1: Different background distributions of genomic features in populations

For the first scenario, we considered the situation that the two training populations differ from each other and both of them are also different from the test population. We also adjusted the simulations to reflect the extend of differences between the two training populations. To simulate such a scenario, we first decided the underlying operational taxonomic unit (OTU) abundance levels for different populations. To do so, we generated three probability vectors to represent the underlying OTU abundance levels in three different populations respectively, adapted from three real colorectal cancer metagenomic datasets. We collected a total of six publicly available and geographically diverse colorectal cancer metagenomic datasets with download links from their original papers [5, 21–25]. We excluded the samples from patients with adenoma so that only samples from patients diagnosed with CRC and healthy controls were used. The numbers of cases and controls as well as country of origin for each dataset are shown in Table 1. We drew a PCoA plot on the samples to show the population differences among these six CRC datasets as in Fig 2. From the PCoA plot, we chose the three least overlapping populations, Hannigan, Feng and Yu datasets as the basis of generating background OTU relative abundance vectors. The Hannigan and Yu datasets were chosen to be used for generating training data, while the Feng dataset was used to generate test data. The two training datasets were pre-processed to retain the top 1000 OTUs with the largest variance in each dataset. We then took the union of the OTUs from the two datasets as the complete OTUs to be used in the following analysis. A total of 1,267 OTUs were used for the simulation study. We kept the OTU of the 1,267 OTUs for the Feng dataset (background distribution for simulating test dataset), and removed the other OTUs. Then, the count data has been transformed to relative abundance vector, using each OTU’s total counts from all samples divided by sum of all OTUs’ total counts.

**Table 1.** Six metagenomic datasets related to colorectal cancer.

| Dataset | Country | No. of cases | No. of controls | Reference |
| --- | --- | --- | --- | --- |
| Zeller | France | 91 | 93 | [21] |
| Yu | China | 74 | 54 | [22] |
| Hannigan | USA/Canada | 27 | 28 | [23] |
| Feng | Austria | 46 | 63 | [24] |
| Vogtmann | USA/Canada | 52 | 52 | [25] |
| Thomas | Italy | 61 | 52 | [5] |

**Fig 2.**
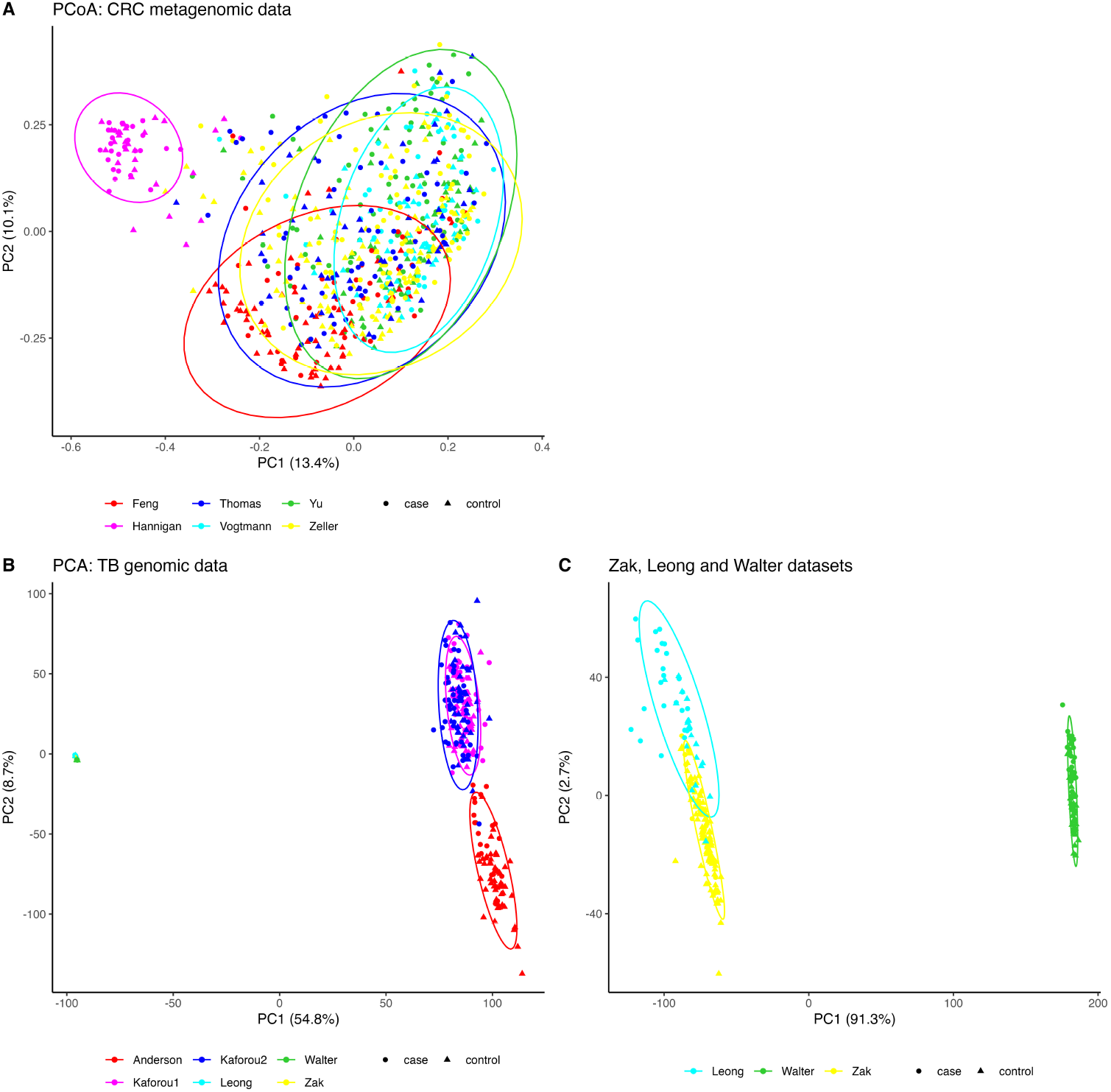
Genomic feature distributions from multiple colorectal cancer and tuberculosis studies. A: Principal coordinate analysis of Bray–Curtis distances computed on six colorectal cancer metagenomic count datasets. B: Principal component analysis of on six tuberculosis gene expression datasets. PCA was computed on the logarithm of fragments per kilobase of transcript per million mapped reads (log FPKM). Zak, Leong and Walter datasets are overlapped in this figure and far away from the other three datasets. C: Principal component analysis of on Zak, Leong and Walter tuberculosis gene expression datasets. Ellipses represent the 95% confidence level assuming a multivariate t-distribution. Round dot represents a case sample, while triangle dot represents a control sample.

Let *v*_1_, *v*_2_, and *v*_3_ be the three background relative abundance vectors from the microbial abundance profiles of the healthy control samples in the pre-processed Hannigan, Yu and Feng datasets, respectively. The three vectors have the same dimension of 1,267, with each dimension as relative abundance level for each OTU. To investigate the impact of difference between populations on cross-study prediction, we created a pseudo-population with relative abundance vector.

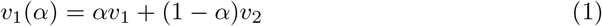

Note that *v*_1_(*α*) − *v*_2_ = *α* (*v*_1_ − *v*_2_). Therefore, the difference between the two simulated populations based on *v*_1_(*α*) and *v*_2_ increases with *α*. When *α* = 0, the two simulated populations have the same underlying distribution and thus there are no population differences between the two training populations. When *α* = 1, the two simulated populations have the largest difference. We used different values of *α* from 0 to 1 with 0.2 increment to represent different magnitudes of training population differences in the following analysis. The relative abundance profiles *v*_1_(*α*) and *v*_2_ were used as background relative abundance vectors for training populations while *v*_3_ was used for the test population.

From these 1,267 OTUs, we randomly chose 10 OTUs and assumed these OTUs were associated with a particular disease of interest. Since disease associated OTUs can be either enriched or depleted, we assumed the first 5 OTUs to be enriched and the other 5 to be depleted. These 10 OTUs were fixed for all the experiments in following analysis. In order to quantify the disease effect on those disease associated OTUs, we defined a disease effect factor, *e*_*d*_, and assumed that the relative abundance of those OTUs were as follows,

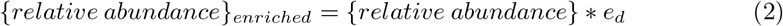

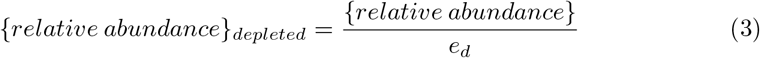

We first modified *v*_1_(*α*), *v*_2_, and *v*_3_ this way and then normalized them to be probability vectors denoted as 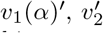, and 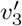, respectively. We then used 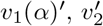, and 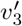 to simulate case microbiome profiles. To see the impact of disease effect on the results, we let *e*_*d*_ to be 1.05, 1.075 and 1.1 in our simulation studies. The larger the value of *e*_*d*_ is, the more marked difference between case and control samples is.

We simulated the OTU counts for the controls and cases in each population as follows. We used one million reads as the library size for all the following simulations. We generated the OTU counts using multinomial distribution *MN* (*library*_*size*_, *v*) where *v* is the relative abundance vector. When simulating the control samples, *v*_1_(*α*), *v*_2_, and *v*_3_ were used in multinomial distribution; when simulating case samples, 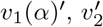, and 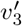were then used. Fifty controls and fifty cases were generated for each *v*, and we referred to the generated datasets as *training*1, *training*2, and *test*, respectively. Then, we changed the count data to log-transformed relative abundance data. The count data were changed to relative abundance data by dividing the sample counts first, followed by a zero-replacement strategy suggested by Martn-Fernndez et al. [26]. In the strategy, we took the minimum non-zero abundance in the dataset, and replaced all the 0 abundances with 0.65 times this minimum non-zero abundance. Then, the non-zero relative abundance data were log-transformed and used in all of the following analysis.

#### Scenario 2: Different batch effects in studies with the same background distribution of genomic features in a population

For this simulation, we kept the underlying populations the same for the training and test data. The underlying OTU abundance profiles were chosen from Yu et al. [22] shown in Table 1. Fixing the background genomic distribution ensures there is no population variation described in Scenario 1 between the training and test samples. The number of disease associated OTUs was set to 10 and library size is one million reads, while the disease effect factor *e*_*d*_ was set to 1.025. We generated 50 cases and 50 controls for each of the two training datasets and one test dataset.

We followed similar procedures as in Zhang et al. [8] to simulate batch effects on the training data. The two training datasets were used as two batches to simulate different batch effects. The batch effect generating model was based on the linear model proposed in ComBat batch correction method [12], which assumes an additive effect on the mean of normalized OTU abundances, and a multiplicative effect on the variance. We then chose three severity levels of batch effects (the levels of changes) for the effect on the mean *sev*_*mean*_ ∈ {0, 3, 5} and three severity levels for the effect on the variance *sev*_*var*_ ∈ {1, 2, 4}. Thus, the model generated batch effect on the two batches with adjusted mean to be {*mean* − *sev*_*mean*_, *mean* + *sev*_*mean*_}, and adjusted variance {*var/sev*_*var*_, *var*× *sev*_*var*_}. Batch effects were only simulated on the training data while the test dataset was unchanged. The two training batches and one test dataset were named as *batch*1, *batch*2, and *test*, respectively. *batch*1 and *batch*2 served the same purpose as *training*1 and *training*2 from the previous scenario.

#### Scenario 3: Different disease models in different studies

In the simulation of different diseased models, unlike the previous two scenarios where 10 disease associated OTUs were fixed in the training and test datasets, we tuned an additional parameter ‘overlapping OTUs’ in the test dataset. The 10 disease associated OTUs were fixed in training data, while the number of disease associated OTUs in test data were chosen from 2, 4, 6, 8, and 10 among the 10 disease associated OTUs in the training data. As the number of overlapping disease associated OTUs increases, the disease models in the training and test data becomes more similar. When the number of overlapping OTUs achieves 10 in the test data, the disease models in the training and test data are the same. After choosing the overlapping OTUs between the training and test disease models, we followed the same procedures as the previous two scenarios to simulate two training datasets with background OTU distribution from the CRC dataset by Yu et al. [22], and one test dataset with background OTUs distribution from the dataset by Feng et al. [24]. The details about the two datasets are shown in Table 1. The three datasets were simulated with the following parameters: 100 samples with 50 cases and 50 controls, one million reads, disease effect factor *e*_*d*_ 1.075. The two training datasets and one test dataset were named as *training*1, *training*2, and *test*, respectively. No batch effects were added in this simulation.

### The random forests (RF) classifiers

After we simulated two training datasets and one test dataset as described above, we treated *training*1 and *training*2 (*batch*1 and *batch*2 in scenario 2) as the training datasets to be used in training RF classifier, and *test* to be used in applying trained classifiers. We randomly split *test* into 50% validation data and 50% test data, and renamed them *val* and *test*. The case and control samples were split evenly to avoid bias. We then pooled the *training*1 and *training*2 together to form a *merged* dataset as suggested by the merging method. *training*1 and *training*2 each has 100 samples, and *val* and *test* each has 50 samples, while *merged* has 200 samples in total. All the RF classifiers trained in our study were implemented by the ‘caret’ package in R [27] with 1000 decision trees, and the ‘mtry’ parameter was tuned by a 10 fold cross-validation. We also chose to use ‘ranger’ as the method for the train function, as it reduced the running time compared to the ‘rf’ method.

### Naive and ComBat normalization settings

We implemented two settings for the experiment, naive and ComBat normalization. In the naive setting, the RF classifiers were trained directly on the original training datasets and applied to the test dataset, and then combined by the integration methods. In the ComBat normalization setting, the *test* dataset was used as reference to normalize *training*1 and *training*2 by ComBat separately. Then, *training*1 *combat* and *training*2 *combat* was generated after the normalization while the *test* was not changed. *merged combat* was generated by pooling *training*1 *combat* and *training*2 *combat* into one dataset and used to be compared with *merged* dataset. RF classifiers were trained on those normalized datasets as well. The prediction results without integration methods using the above RF classifiers are described as following.

1. **Training1:** Train RF classifier on *training*1.
2. **Training2:** Train RF classifier on *training*2.
3. **Training1 ComBat:** Train RF classifier on *training*1 *combat*.
4. **Training2 ComBat:** Train RF classifier on *training*2 *combat*.
5. **Merged:** Train RF classifier on *merged*.
6. **Merged ComBat:** Train RF classifier on *merged combat*.

For Scenario 2, the notations were slightly different from the above. “Training1”, “Training2”, “Training1 ComBat” and “Training2 ComBat” were renamed as “Batch1”, “Batch2”, “Batch1 ComBat” and “Batch2 ComBat” since we had two batches with different batch effects. “Merged” was replaced by two methods “Merged NoBatch” and “Merged Batch”. “Merged NoBatch” was done by training RF classifier on the original merged dataset with no batch effect simulated; while “Merged Batch” was done by pooling “Batch1” and “Batch2” into one dataset first followed by the trained RF classifier. “Merged ComBat” was done by pooling “Batch1 ComBat” and “Batch2 ComBat” into one dataset, followed by the trained RF classifier. The above trained classifiers were then applied to test data as predictors. Then, different ensemble weighted learning or rank aggregation methods combined the predictors from “Training1” and “Training2” or from “Training1 ComBat” and “Training2 ComBat” to get final predictor.

### Ensemble weighted learning methods

Patil et al. [9] evaluated the performance of cross-study learner by using five alternative choices of weights: simple average of predictions from each single-study learner (“Avg”), average weighted by study sample size (“n-Avg”), average weighted by cross-study performance (“CS-Avg”), stacked regression (“Reg-s”), and averages of study-specific regression weights (“Reg-a”). The last three ensemble methods reward reproducibility. We compared these 5 ensemble weighted learning methods in our study as well. In addition, we also implemented two other methods combined with RF classifier as implemented in [13]. The detailed ensemble weighted learning methods are described as follows.

#### 1. Avg

Take the average of predictors from “Training1” and “Training2”.

#### 2. n_Avg

Average weighted by study sample size of the predictors from “Training1” and “Training2”. In the simulations, since the sample sizes are the same for the two training datasets, “n Avg” is the same as “Avg”. For real data applications, due to different sample sizes of the datasets, the results from these two methods can be different.

#### 3. CS-Avg

Average weighted by cross-study performance. RF classifier was trained on *training*1 and the applied the trained classifier on the samples from *training*2 to get predicted probabilities, then calculated the cross-entropy loss. Cross-entropy loss is defined as:

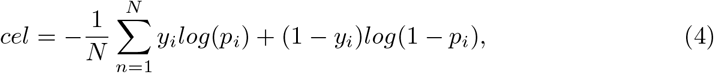

where *N* is the sample size in the test dataset, *y*_*i*_ is the real case/control status of sample *i* (*y*_*i*_ is 0 if it is a control sample, or is 1 if it is a case sample), and *p*_*i*_ is the predicted probability of sample *i* to be a case sample by the machine learning classifier.

The cross-entropy loss of training on *training*1 and predicted on *training*2 was calculated and named as *cel*1. Likewise, we trained RF classifier on *training*2 and predicted on *training*1, and calculated the cross-entropy loss *cel*2. Next, calculated weights as *weight*1 = | *cel*1 − *max*(*cel*1, *cel*2|), and *weight*2 = |*cel*− 2 *max*(*cel*1, *cel*2) |, and normalize the two weights to have sum of 1. In the two training datasets scenario, the method always assigns the RF classifier with worse cross-study performance (higher cross-entropy loss) to have zero weight, so in the simulations, only one of the predictors from “Training1” and “Training2” was used for this method. In general scenario where multiple training datasets are presented, the cross-entropy loss is calculated for each training dataset performing predictions on each of the other datasets, for example, *cel*_*ij*_ is calculated by the predicted probabilities from samples in dataset *j* while training on dataset *i*, then for dataset *i* the total cross-entropy loss is calculated by *cel*_*i*_ = ∑_*j,j≠i*_ *cel*_*ij*_. Similar as the two training datasets scenario, the weights for each dataset is calculated as *weight*_*i*_ = |*cel*_*i*_ − *max*(*cel*_1_, *cel*_2_, …, *cel*_*i*_, …, *cel*_*n*_) | where *n* is the total number of training datasets used. The final weights are normalized to sum to 1. This method assigns zero weight to the model with the worst average performances on the rest of the training datasets.

#### 4. Reg-a

Averages of study-specific regression weights computed by non-negative least squares. RF classifiers were trained on *training*1 and *training*2, respectively. Then predicted on *training*1 and *training*2 to obtain 4 predictors. For testing on *training*1, fit non-negative least squares to the two associated probability vectors with real case/control status in *training*1 as response, and get the 2 coefficients. Repeated same procedure for testing on *training*2. After this step, we constructed a 2× 2 coefficient matrix, with each row representing test data and each column representing training data. Then, the coefficients in each row were multiplied by the sample size of the test data, and the weights were finally computed as the column average of the adjusted coefficients.

#### 5. Reg-s

Stacked regression weights computed by non-negative least squares. The 2 predicted probability vectors from the “Reg-a” method were stacked into one vector for each test dataset, and then fit non-negative least squares to the stacked vectors with real case/control status as response. The coefficients were then used as weights.

#### 6. val-auc

RF classifiers were trained on *training*1 and *training*2 independently. Then applied the two trained classifiers on *val* to predict. Calculate the area under the operational characteristic curve (AUC) scores by comparing to the real disease status from *val*, and assign the two AUCs as weights on the predictors from “Training1” and “Training2”.

#### 7. LOSO-auc

Leave-One-Sample-Out (LOSO) AUCs were calculated for all test samples, and then used the corresponding *AUC* 0.5 as weights to combine predictors from “Training1” and “Training2”. This method was proposed in [13].

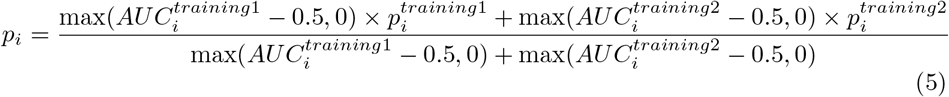

In the ComBat normalization settings, the above seven integration methods were applied similarly to the predictors obtained from “Training1 ComBat”, “Training2 ComBat” and “Merged ComBat”. The area under the operational characteristic curve (AUC) scores were computed by comparing the predicted probabilities from the combined predictors by ensemble learning methods to the real case/control status, and evaluated the performance of each of these methods.

### Rank aggregation methods

Rank aggregation methods have not been used for predicting phenotypes. In this study, we investigate how to use rank aggregation for phenotype prediction and their performance. For each individual prediction method, we first ranked the samples in the test data based on the predicted probabilities of being cases in a decreasing order to obtain a ranked list, and assigned the smallest rank to items with ties. We then created two ranked lists based on the predictions from the two training predictors. Next we applied some rank aggregation methods to obtain an aggregated rank list for the samples.

We used five rank aggregation methods and compared their performance both among themselves and to the previous mentioned ensemble learning methods.

#### 1. mean

Take the mean of the two rank lists.

#### 2. geometric mean

Take the geometric mean of the two rank lists.

#### 3. Stuart rank aggregation

First, normalize the ranks to rank ratios. Then, use order statistics as proposed by Stuart et al. [28] to create an aggregated rank list. We used the ‘RobustRankAggreg’ package [29] from R with method ‘stuart’ to compute the results.

#### 4. Robust rank aggregation (RRA)

This method was proposed by Kolde et al. [29] also based on the use of order statistics, but improved on the computational efficiency and statistical stability. For each item in the rank list, the algorithm looks at how it is positioned and compares this to the baseline case where all the preference lists are randomly shuffled. A P-value is then assigned for all items, showing how much better it is positioned in the ranked lists than expected by chance. This P-value is then used to re-rank the list. We used the ‘RobustRankAggreg’ package [29] to compute the results.

#### 5. Bayesian analysis of rank data with covariates (BARC)

Li et al. [30] recently developed a Bayesian based rank aggregation method incorporating information from covariates. Even though covariates are not of concern in our study, we used their rank aggregation method with no covariates involved version. The method is accessible from https://github.com/li-xinran/BayesRankAnalysis

After we obtained the aggregated rank lists by the above methods, we then computed the area under the operational characteristic curve (AUC) scores to evaluate the performance of those rank aggregation methods. Similar to the ensemble weighted learning methods, those rank aggregation methods were applied to both naive and ComBat normalization settings separately.

### Applications on real CRC metagenomic datasets

We demonstrated applications of merging and integration methods to six real CRC metagenomic datasets. We excluded the samples from patients with adenoma so that only samples from patients diagnosed with CRC and healthy controls were used. The information about the numbers of cases and controls for each dataset, country of origin, and relevant references are shown in Table 1. We used MicroPro [31] to generate microbial count profiles for the six CRC datasets. The generated count data has been log transformed into relative abundance data using the same procedures as described in the data pre-processing in simulation studies Scenario 1: Different background distributions of genomic features in populations. We then pre-processed each dataset to retain the top 1000 OTUs with the largest variance.

We implemented a Leave-One-Dataset-Out (LODO) setting for the real data applications. For each dataset among the six datasets, we treated it as a test data and the other five datasets were used as training data. We took the union of all the OTUs in the five training datasets and filled with 0 abundance when the feature was missing in any particular dataset. We applied ComBat normalization on the five training datasets as in simulation studies, while the training data were not changed for the naive setting. Next, we trained RF classifiers on the five training datasets independently, and applied integration methods to the five predictors. For the merging method, the five training datasets were pooled into one dataset, and a single RF classifier was trained on it.

### Applications on TB gene expression datasets

For the real application on the TB gene expression studies, we collected six annotated and cleaned datasets from Zhang et al. [8]. The information about the country of origin, numbers of case and controls, and relevant references can be found in Table 2. The TB datasets were already transformed into the logarithm of fragments per kilobase of transcript per million mapped reads (logFPKMs). Therefore we directly selected the top 1000 gene features with the largest variance for each of the datasets, and took the union of those genes as the complete features to be used in the following analysis. The Leave-One-Dataset-Out strategy was performed on the six TB gene expression datasets in the same way as for the CRC datasets described above Applications on real CRC metagenomic datasets.

**Table 2.** Six gene expression datasets related to tuberculosis.

| Dataset | Country | No. of cases | No. of controls | Reference |
| --- | --- | --- | --- | --- |
| Zak | South Africa/Gambia | 16 | 104 | [32] |
| Anderson | South Africa/Malawi/Kenya | 20 | 50 | [33] |
| Leong | India | 25 | 19 | [3] |
| Walter | USA | 35 | 35 | [34] |
| Kaforou1 | South Africa | 46 | 48 | [35] |
| Kaforou2 | Malawi | 51 | 35 | [35] |

## Results

### ComBat normalization is essential for heterogeneous populations

In the first scenario, we investigated the impacts of the different background operational taxonomic unit (OTU) distributions in the training samples on the prediction performance and the results are shown in Fig 3. When the training and the test data have different background OTU distributions, direct applications of the trained prediction model based on the training data to the test data yielded very low prediction accuracy with area under the operational characteristic curve (AUC) close to 0.56 as the “Training1” and “Training2” rows show. Merging the raw training data and directly integrating the trained models from multiple training samples did not improve the prediction accuracy. These results clearly showed the importance of normalization in building prediction models.

**Fig 3.**
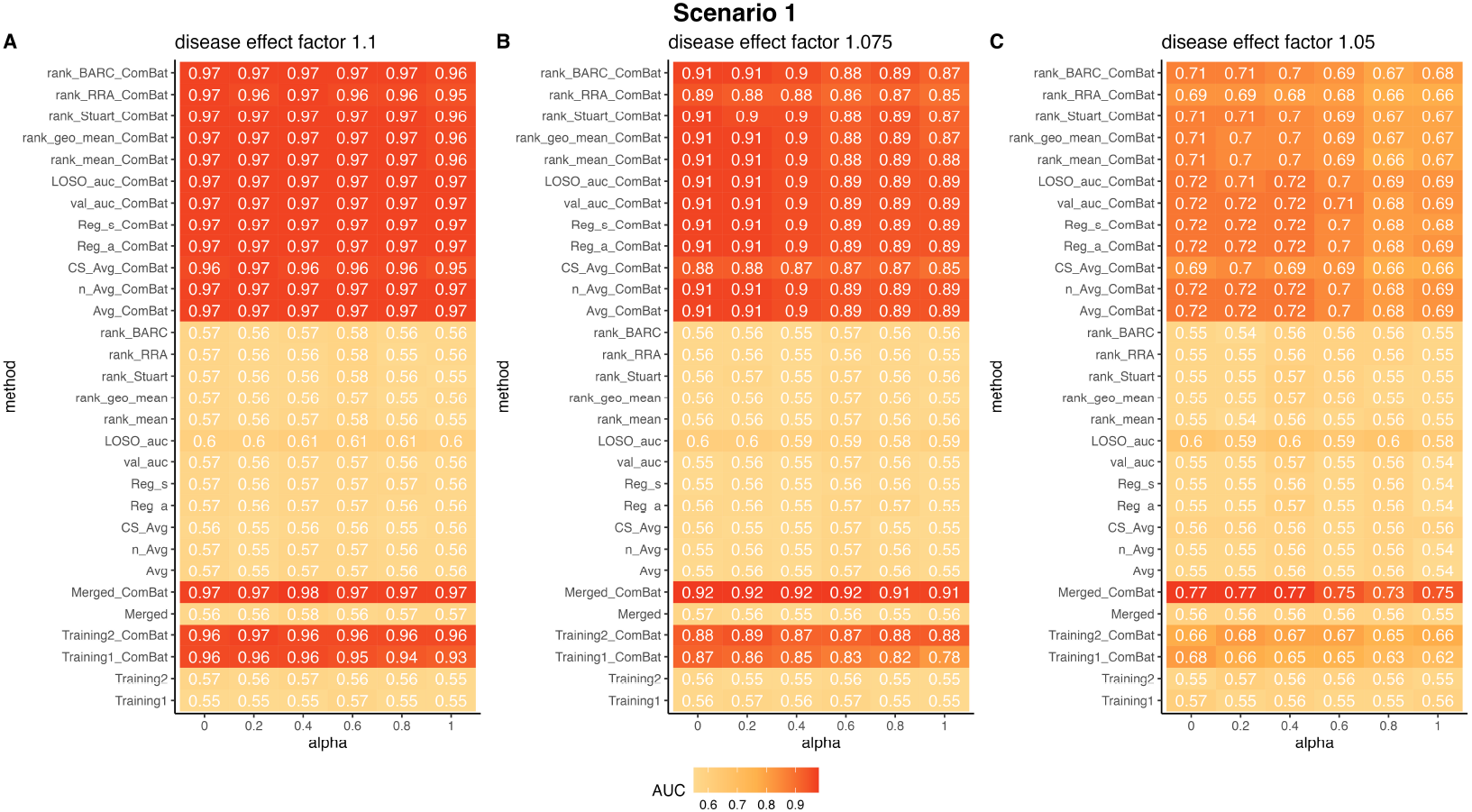
ComBat normalization markedly increases cross-study prediction when the training and test data have different feature distributions. The figures show the AUCs of RF prediction using different integration methods with three different disease effect factors. Columns represents different values of *α*. All the method names without a suffix of “ComBat” are the methods carried out in the naive setting, while the names with a suffix of “ComBat” were carried out in the ComBat normalization setting. All the experiments were repeated for 100 times and the AUC scores shown on the figure are the averages from the 100 trials.

Many methods have been developed for normalization of metagenomic data across different studies ([12], [15], [16]). However, most of these methods were designed to mitigate the impacts of experimental artifacts, not for dealing with population differences among the different studies. Still we applied one of the widely used normalization method, ComBat [12], to normalize the different metagenomic data from the different populations to see if the normalization can improve the prediction accuracy. It is important to see that using ComBat to normalize the metagenomic data markedly improved the prediction accuracy in the test data. For example, the AUC score for “Training2 ComBat” was increased from average of 0.56 to about 0.96, 0.88 and 0.66 when the disease effect was set at 1.100, 1.075 and 1.050, respectively. For different values of *α*, we used *v*_1_(*α*) = *αv*_1_ + (1− *α*)*v*_2_ as the background OTU relative abundance for “Training1”, where *α* measures the difference between “Training1” and “Training2”. It can be seen that the prediction accuracy decreases as *α* increases for all values of disease effect *e*_*d*_. For example, when *e*_*d*_ = 1.075, the AUC for “Training1 ComBat” decreased from 0.87 to 0.78 as *α* increases from 0 to 1. This observation can potentially be explained by the fact that *v*_1_ is further away from the test data than *v*_2_.

With the marked increase of prediction accuracy for “Training1 ComBat” and “Training2 ComBat”, we investigated if integrating the two predictors by ensemble weighted learning or rank aggregation can further increase the prediction accuracy and the results are shown on the top rows marked with red. Except for “rank RRA ComBat” and “CS Avg ComBat” that slightly underperformed the other integration methods, all the other integration approaches give similar results and they also outperformed both “Training1 ComBat” and “Training2 ComBat”.

Most surprisingly, the relatively simple merging method after ComBat normalization, “Merged ComBat”, yielded the best performance. When the disease effect *e*_*d*_ is relatively high, for example, *e*_*d*_ *>*= 1.075, the increase in AUC over other integration method is minimal. However, when *e*_*d*_ = 1.05, the AUC for “Merged ComBat” is 0.77 compared to about 0.72 for other integration methods when *α* is small. Similarly, the AUC for “Merged ComBat” is 0.75 compared to about 0.69 for other integration methods when *α* is large.

These results indicates normalizing the population difference before training machine learning models could play an important role in improving model prediction performance. This result is surprising given that ComBat was originally designed to correct for experimental artifacts like batch effects, not for adjustment for population differences. Our results clearly showed that ComBat can be used to adjust for population differences to increase cross study prediction accuracy.

### ComBat normalization can successfully remove batch effects within the same population

In the second scenario, we considered studies within the same population but from different laboratories or using different sequencing technologies. In such a scenario, experimental batch effects can happen and it is essential to correct the batch effects in such studies. As presented in Scenario 2: Different batch effects in studies with the same background distribution of genomic features in a population, we simulated batch effects affecting the mean and the variance of OTU abundance levels, respectively. We evaluated the RF classifier’s prediction accuracy on the test data for the two types of batch effects separately. In this scenario, we let disease effect *e*_*d*_ = 1.025 to clearly show the relative performance of the different methods. Fig 4A shows the results when there are additive effects on the mean (*sev*_*mean*_ ∈ {0, 3, 5}) with variance remains unchanged (*sev*_*var*_ = 1). Without data normalization, the AUC score on the test data is slightly higher than 0.5 when *sev*_*mean*_ = 0 and is 0.5 when *sev*_*mean*_ ≠ 0 as expected. After ComBat normalization, the AUC scores on the test data increased to about 0.75 for all the parameter values. Most of the ensemble weighted learning and rank aggregation methods of the two predictors further increased the AUC close to 0.8. Similar to scenario 1, “rank RRA ComBat” and “CS Avg ComBat” slightly underperformed other integration methods. Simply merging the two training datasets after ComBat normalization achieved the highest AUC score of about 0.85. The last row showed the highest AUC score one could get if there were no batch effects and it shows that the prediction accuracy was only slightly decreased from 0.88 to 0.85 with “Merged ComBat”.

**Fig 4.**
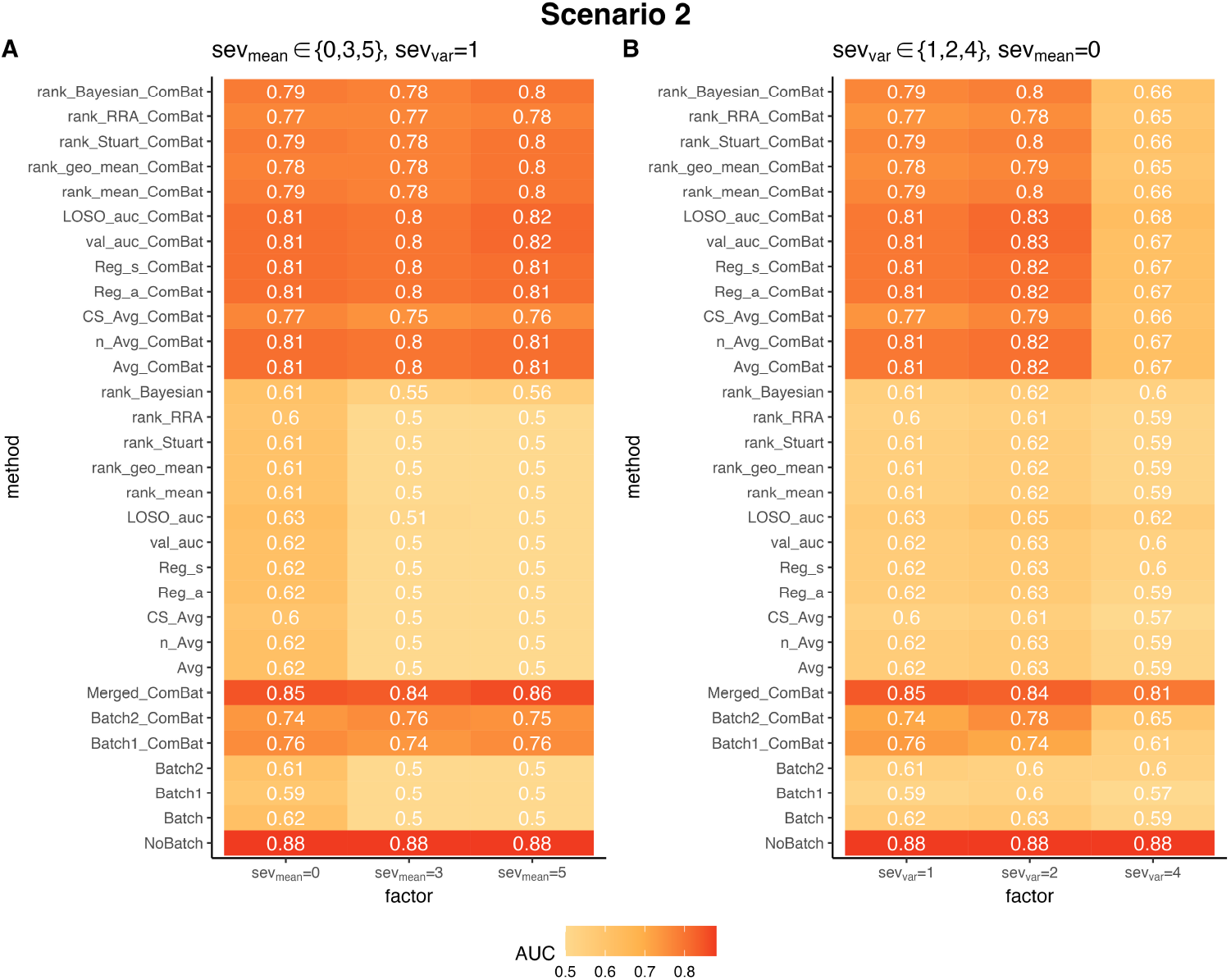
ComBat normalization markedly increases cross-study prediction when the studies have batch effects. The figures show the AUCs of RF prediction using different integration methods with various batch severity levels. A: AUC score comparisons with different severity levels of additive batch effects on the mean of OTU abundances, with no multiplicative batch effect on the variance. B: AUC score comparisons with different severity levels of multiplicative batch effects on the variance of OTU abundances, with no additive batch effect on the mean. The disease effect factor was set to 1.025 for both situations. All the method names without a suffix of “ComBat” are the methods done in naive setting, while the names with a suffix of “ComBat” were done in ComBat normalization setting. All the experiments were repeated for 100 times and the AUC scores shown on the figure are the averages from the 100 trials.

Fig 4B shows the results when there are multiplicative effects on the variance (*sev*_*var*_∈ {1, 2, 4}) with no effect on the mean of OTU abundance levels (*sev*_*mean*_ = 0). Since the mean perturbation is 0, without normalization there are still some predictive powers for the test data although the AUC score is generally low at around 0.6 based on the two batches separately. Integrating the two predictors without ComBat normalization did not improve the prediction accuracy. When *sev*_*var*_ = 1 or 2, after ComBat normalization, the AUC scores based on the two training datasets were increased to about 0.75. Integrating the predictors after ComBat normalization further increased the AUC to about 0.80. On the other hand, when *sev*_*var*_ = 4, the ComBat normalization only increased the AUC to 0.67 using ensemble weighted learning and rank aggregation. However, merging after ComBat normalization, “Merged-ComBat”, achieved the highest AUC score of 0.81.

### Prediction accuracy can be markedly decreased as the number of overlapping disease associated OTUs decreases

In Scenario 3, we investigated how the differences of disease models in the training and test data affect the RF classifier’s prediction AUC scores and the results are shown in Fig 5. The relative performances of the different prediction methods were mostly consistent with the results from the first two scenarios. However, we did not see obvious advantage of “Merged ComBat” over other ensemble weighted learning and rank aggregation methods although “Merged ComBat” was still one of the best performers.

**Fig 5.**
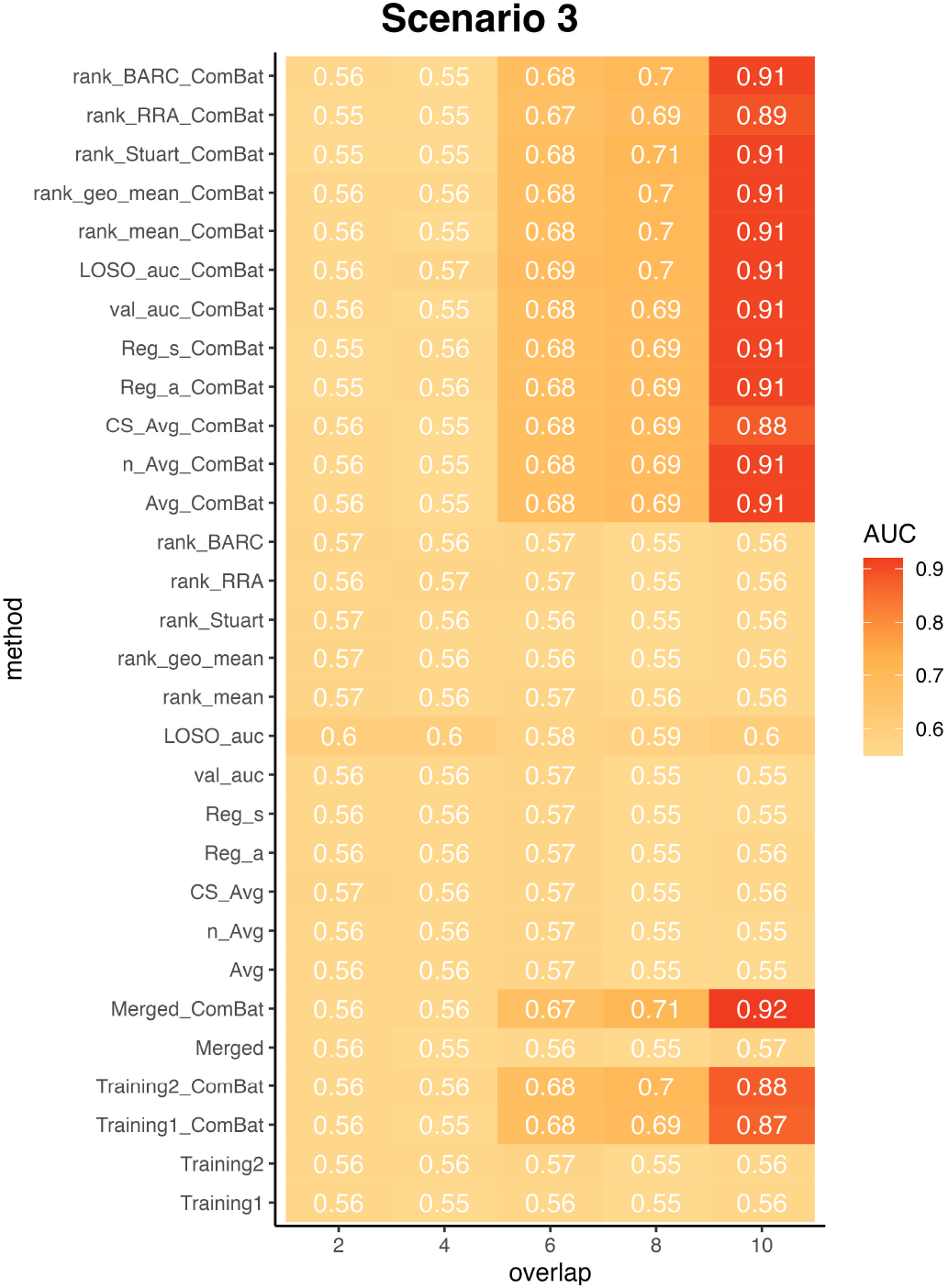
ComBat normalization markedly increases cross-study prediction when the training and test data have different disease models. The figures show the AUCs of RF prediction using different integration methods with various number of overlapping disease associated OTUs. The disease effect factor was set to 1.075. Columns represent different numbers of overlapping disease associated OTUs in the training and test data, the larger the number, the more similar the two disease models are. When the number achieves 10, the two models are the same in the training and test data. All the experiments were repeated for 100 times and the AUC scores shown on the figure are the averages from the 100 trials.

As expected, the prediction accuracy is markedly decreased as the number of overlapping disease associated OTUs decreases. For example, when *e*_*d*_ = 1.075 with only less than 4 overlapping associated OTUs between the training and test data, the AUC is less than 0.57 on average. When the number of overlapping disease associated OTUs increases to 6 to 8, the optimal AUCs increased to about 0.7. On the other hand, when the number of overlapping disease associated OTUs increased to 10, the optimal AUC is around 0.91.

### Aggregating ranks from predicted probabilities is an alternative powerful way to integrating multiple heterogeneous studies

Aggregating the rank lists of the genes from multiple studies in order to have a complete understanding of the biological interest has been frequently used in genomic studies. Rank aggregation provides insights to integrating heterogeneous studies without dealing with the challenge of normalizing the data across these studies, and previous studies have shown that the aggregated rank list provides more meaningful results than single rank list ([28–30]). In our study, instead of aggregating the ranks of the genomic features from multiple studies, we developed a method integrating the predicted probabilities of test samples from multiple machine learning classifiers by transforming those probability lists into rank lists, followed by different rank aggregation methods. In our study, we focused on five established rank aggregation methods: mean of ranks, geometric mean of ranks, Stuart rank aggregation [28], Robust Rank Aggregation (RRA) [29] and Bayesian Analysis of Rank data with Covariates (BARC) [30]. The detailed descriptions of these methods are given in Rank aggregation methods.

In our simulation studies, we observed similar prediction performance of the five rank aggregation methods to the ensemble weighted learning methods in all three scenarios (Fig 3, Fig 4 and Fig 5). In particular, all these five rank aggregation methods performed well when combined with ComBat normalization.

In the real applications of colorectal cancer (CRC) and tuberculosis (TB), the five rank aggregation methods showed slightly lower prediction performance than the ensemble weighted learning methods under naive settings. However, when combined with ComBat normalization, all the five methods achieved similar and promising prediction results compared with ensemble weighted learning methods. Surprisingly, S1 Fig and S2 Fig show that for individual studies, in the cases when ensemble weighted learning didn’t improve the AUC scores (Figure S1 B and D), the rank aggregation methods actually improved the prediction performance when compared with the ensemble weighted learning methods.

As shown in both simulations and real applications, aggregating the ranks of samples’ predicted probabilities provides new insights when integrating multiple heterogeneous datasets in terms of phenotype prediction. We demonstrated that the rank aggregation methods are as robust as the ensemble weighted learning methods, and can even boost the prediction performance in some cases when ensemble weighted learning didn’t work well.

### Applications to metagenomic datasets related to colorectal cancer

We analyzed 6 metagenomic datasets related to colorectal cancer (CRC) as summarised in Table 1 using the various methods. We implemented a Leave-One-Dataset-Out (LODO) setting for the analyses with one of the six datasets as test data, while the other five datasets as training data. The detailed method is described in Applications on real CRC metagenomic datasets. Fig 6A shows the average AUCs among the six LODO experiments for the “Merged”, “Ensemble weighting”, “Ensemble weighting (normalized)”, “Rank aggregation” and “Rank aggregation (normalized)”. The “Single learner” results from each LODO experiment were the averages of 5 single learners (5 training datasets), followed by the average among the six LODO experiments. The independent results for each LODO experiment can be found in S1 Fig.

**Fig 6.**
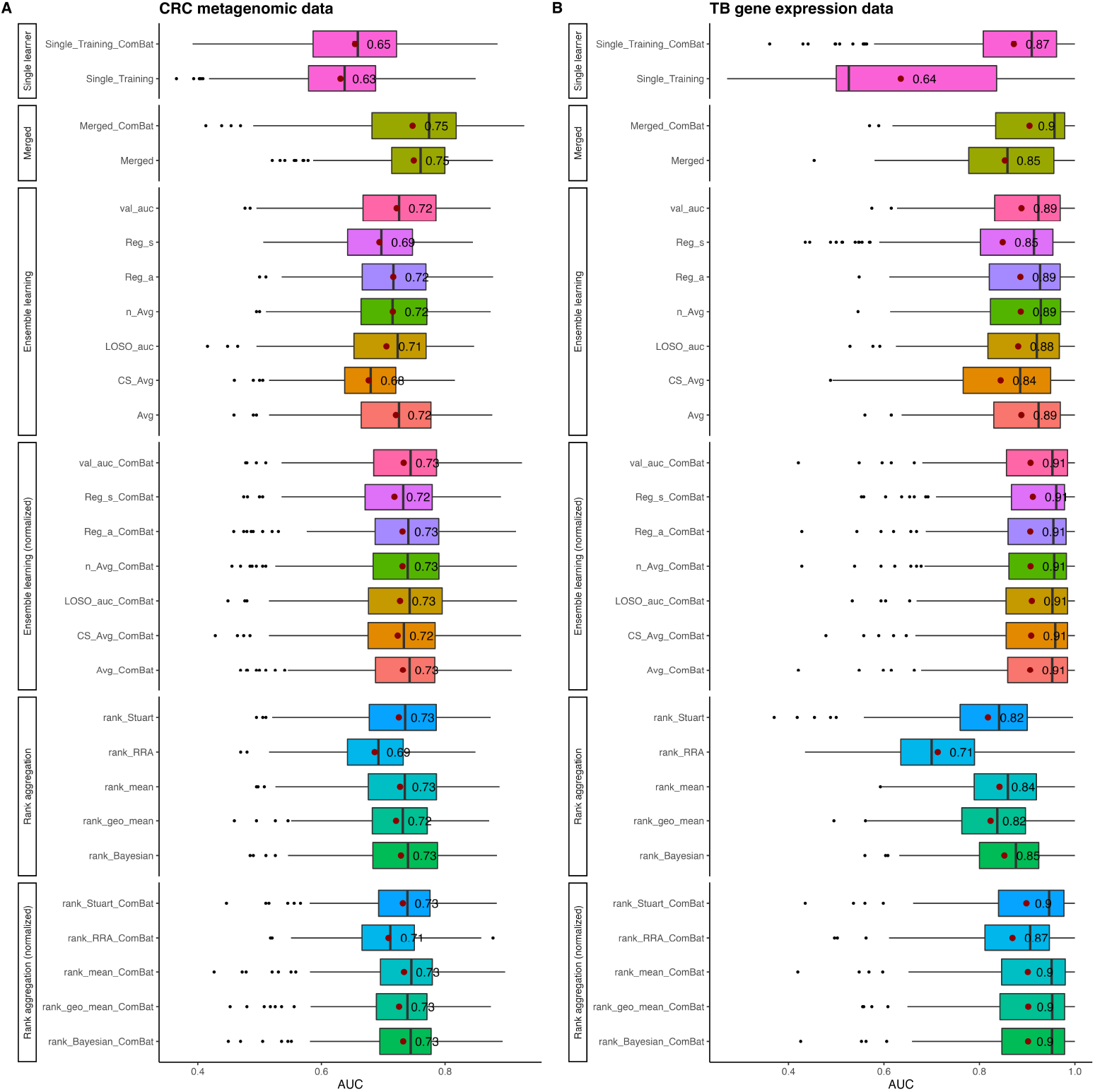
Realistic applications of merging and integration methods on multiple colorectal cancer metagenomic datasets and tuberculosis gene expression datasets. A: Leave-one-dataset-out average AUC score comparisons among different methods in colorectal cancer metagenomic datasets. B: Leave-one-dataset-out average AUC score comparisons among different methods in tuberculosis gene expression datasets. The results by different methods are grouped into six groups. “Single learner”: Each of the five training datasets were trained independently with RF classifier and predicted on the test dataset, then the average AUC score was taken among the five predictions. “Merged”: Merging method with pooling all five training datasets into one train data. The “Single learner” and “Merged” experiments were conducted under both naive and ComBat normalization settings. “Ensemble learning”: The five training predictors were integrated by ensemble weighted learning methods under naive setting. “Ensemble learning (normalized)”: The five training predictors were integrated by ensemble weighted learning methods under ComBat normalization setting. “Rank aggregation”: The five training predictors were integrated by rank aggregation methods under naive setting. “Rank aggregation (normalized)”: The five training predictors were integrated by rank aggregation methods under ComBat normalization setting. The red dots and associated values on the figure are the mean AUC scores for each method, while the black dots represent the outliers. Same method under different settings are represented in the same color of boxplots. All the experiments were repeated 30 times for each test dataset, and the results presented in the figure were based on the average AUC scores of the total 180 replications for the six test datasets for metagenomic and gene expression studies respectively.

As shown in Fig 6A, when one single training dataset was used and predicted on the test data, the AUC result increased from 0.63 to 0.65 on average when we used ComBat normalization on the training data. The AUC results for ensemble weighted learning and rank aggregation both improved slightly after ComBat normalization, but none of those integration methods produced better AUC results compared with merging the five training datasets. Comparing the prediction performance from the merging and integration methods with the single learning, it is expected that the AUCs increased about 0.1 on average, and this further supports the idea that cross-study prediction using multiple training datasets is more accurate than using single training dataset.

From the individual training results as in S1 Fig, we observed similar trends as in Fig 6A in most cases. Among the six test datasets, the AUCs associated with the Hannigan dataset has the lowest scores, and neither merging with ComBat normalization or ensemble weighted learning methods improved the prediction performance. Also, the AUC scores when using Hannigan as test data is the lowest among the six results, with an average of 0.61 for single learner and 0.63 for integration methods. This observation is consistent with the data distribution from the PCoA plot Fig 2A. As shown on the figure, the Hannigan dataset is most separate dataset from the other five datasets with least overlap, and this demonstrated the very different distribution of the count data from the Hannigan dataset than the other five datasets. As we illustrated by the simulation studies, different background distributions of genomic features in populations compromised the reproducibility of machine learning classifier’s prediction performance.

### Applications to gene expression datasets related to tuberculosis

To further investigate the prediction performance of merging and integration methods on real datasets, we used 6 tuberculosis (TB) gene expression datasets as summarised in Table 2, and repeated the procedures in Applications to metagenomic datasets related to colorectal cancer to the gene expression data.

In the application of TB studies (Fig 6B), the overall AUC results were much higher than the CRC studies, and the ComBat normalization results improved markedly on average among all analyzed methods. Unlike the application of CRC studies, the ensemble weighted learning methods outperformed slightly the merging method in both naive and ComBat normalization settings, while the rank aggregation method performed slightly better than the merging method under the ComBat normalization setting.

From the individual plots S2 Fig, we observed that when the Zak and Anderson datasets were served as test data, the prediction results are lower than the other four datasets, and the ComBat normalization improvements on all methods are smaller than the other four as well. This observation is consistent with the study by Zhang et al. [8] where these two datasets has highest cross-entropy losses when served as test data. The possible explanation for these difference is that the two datasets include only children or adolescents, while the other four datasets include adults as well. According to Alcaïs et al. [36], children and adults developed different tuberculosis clinical features and parthenogenesis, and this could affect the machine learning models’ reproducibility when the populations have different features in terms of disease.

We also observed the remarkable differences among training different single learners. For example, in Figure S2 A, when RF classifier was trained on the Walter and Leong datasets and predicted on the Zak dataset, the AUC results are much higher than trained on Anderson, Kaforou1 and Kaforou2. Similarly, when training RF classifier on Kaforou1 and Kaforou2 and predicted on Anderson, the AUCs can achieve 0.83 and 0.88, respectively, while training on the other three datasets and predicted on Anderson only has 0.5 AUC which is more like a random guess. These observations are consistent from the data distribution in the PCA plots (Fig 2B), Anderson, Kaforou1 and Kaforou2 are closer to each other and far away from Zak, Walter and Leong, while the later three datasets are closer to each other. This further indicates that the heterogeneity in different datasets has large impacts on the reproducibility of machine learning classifiers.

Lastly, based on the results from the real applications on the CRC and TB studies, consistent with the results of simulation studies mentioned above, we demonstrated that the ComBat normalization method is essential for heterogeneous populations. When dealing with heterogeneous populations, it is recommended that using both merging and integration methods to find best prediction results.

## Discussion

With the increasing availability of large collections of omic data, the reproducibility of machine learning prediction models has raised great concerns when conducting cross-study predictions with the impact of study heterogeneity. Previous studies have addressed this issue and developed many statistical methods to overcome study heterogeneity, including merging with batch effect removal [37] and ensemble learning methods [9]. In this study, we performed a comprehensive analysis of different methods on the phenotype prediction by integrating heterogeneous omic studies. We considered three different sources of heterogeneity between datasets, including population differences, batch effects and different disease models. We developed a workflow in simulating these three sources of heterogeneity and generating simulated samples based on real datasets. We also evaluated the prediction performance of many different statistical methods, including merging, ensemble weighted learnings and rank aggregations. Besides the comparisons of different methods, we also explored the potential of normalizing the data by ComBat first then applied those statistical methods mentioned above. We provided both simulation studies and real data applications on CRC metageomic datasets and TB gene expression datasets to compare different approaches.

In our simulation studies, we observed a decreasing trend in prediction accuracy among all statistical methods we investigated when the population heterogeneity became large, the batch effects increased on training data, as well as the differences of disease models between training and test data enlarged. These observations indicate that overcoming the heterogeneity needs to be addressed before applying machine learning prediction models on cross-study settings. Merging and integration methods that integrate different studies for phenotype predictions without batch correction did not improve the prediction accuracy much when compared to single training model, but when combined with ComBat normalization, we observed a remarkable improvement in the the prediction accuracy in all simulations. These observations indicate normalizing the heterogeneous datasets before training machine learning models is essential in improving phenotype prediction performance.

It is noteworthy that our simulations yielded different conclusions in contrast with the study by Zhang et al. [8] on the prediction performance of using merging with ComBat normalization methods. In their study, they showed that merging with ComBat normalization was not as robust as ensemble learning methods at high severity of batch effects. However, in our second scenario of simulating different severity of batch effects on training and test data, we observed that merging combined with ComBat normalization always achieved highest prediction performance in spite of severity of batch effects (Scenario 2: Different batch effects in studies with the same background distribution of genomic features in a population). We investigated the contradictions of observations in our study and the study by Zhang et al. [8], and we noticed that the ComBat normalization process in our study was different from theirs. In their study, when multiple training batches were provided, the batches were pooled into one training data, then applied ComBat normalization on the pooled dataset. They then did a second round of ComBat normalization on the test data using the pooled training data as reference. In our study, we normalized the training batches using the test data as reference when conducting ComBat normalization independently. The normalized training batches were pooled into one data for training machine learning model. Since test data is always the target of prediction, instead of adjusting the test data, we used it as the baseline to adjust different training batches so that the differences between training and test data were mitigated more effectively. Our study showed that our normalization approach yielded higher prediction accuracy.

Consistent with simulations, the applications on the CRC metagenomic and TB gene expression datasets with Leave-One-Dataset-Out experiments showed similar trends in terms of performance of merging and integration methods combined with ComBat normalization. However, the increasing trend in prediction accuracy is less remarkable than in the simulation studies, probably caused by the escalations of heterogeneity when five training datasets were used. Different to the simulations, when the number of training datasets became five in the real data applications instead of two in the simulations, the prediction performance of all merging and integration methods improved compared to prediction with single training dataset even without ComBat normalization. These results also suggest that the importance of integrating multiple studies than only using single study in terms of machine learning model reproducibility.

With the comparisons of the statistical methods used in our study, we saw similar trends for all the ensemble weighted learning methods except the slightly lower performance of the “CS-Avg” method. The “CS-Avg” method penalized the training dataset with the worst average performances on the rest of the training datasets when doing cross-training-data validations, and it excludes this dataset from predicting on test dataset. The worse performance of “CS-Avg” demonstrated that excluding the worst performance training data may not be beneficial to phenotype predictions as it discards useful information from that particular training data in the same time.

Therefore, we suggest to use other ensemble weighted learning methods that also penalize worst performance training data but retain the useful information in some way. We also incorporated the rank aggregation methods into our study as well, and illustrated that the rank aggregation methods showed similar prediction performances, which also boosted the prediction accuracy remarkably. Rank aggregation methods should be considered as an alternative way for integrating heterogeneous studies in the future. We also noticed the extraordinary performance in merging method, and consistent with the findings by Guan et al. [38], merging and integration methods can outperform each other in different scenarios, and when training multiple studies we should consider to use both methods to find optimal.

## Acknowledgements

We thank Beibei Wang for advice on choosing and implementing batch normalization method.

## Availability of data and materials

All the CRC datasets used in this study are publicly available in the European Nucleotide Archive (ENA) database (https://www.ebi.ac.uk/ena). Accession number for Yu is PRJEB10878 [22], for Hannigan is PRJNA389927 [23], for Feng is ERP008729 [24], for Vogtmann is PRJEB12449 [25], for Zeller is ERP005534 [21], and for Thomas is SRP136711 [5]. All the TB annotated and cleaned datasets used in this study are obtained from the study by Zhang et al. [8] and available on their github repository: https://github.com/zhangyuqing/bea_ensemble All the codes used in analysis can be found at https://github.com/lynngao/Heterogeneous-Studies

## Author Contributions

F.S. conceived and designed the study, and supervised the work. Y.G. implemented the methods, conducted the computational analyses, and drafted the manuscripts. F.S. modified and finalized the manuscripts. All authors agree to the content of the final paper.

## Supporting information

**S1 Fig. Realistic applications of merging and integration methods on multiple colorectal cancer metagenomic datasets**. The results by different methods are grouped into six groups. “Single learner”: Each of the five training datasets were trained independently with RF classifier and predicted on the test dataset, then the average AUC score was taken among the five predictions. “Merged”: Merging method with pooling all five training datasets into one train data. The “Single learner” and “Merged” experiments were conducted under both naive and ComBat normalization settings. “Ensemble learning”: The five training predictors were integrated by ensemble weighted learning methods under naive setting. “Ensemble learning (normalized)”: The five training predictors were integrated by ensemble weighted learning methods under ComBat normalization setting. “Rank aggregation”: The five training predictors were integrated by rank aggregation methods under na ïve setting. “Rank aggregation (normalized)”: The five training predictors were integrated by rank aggregation methods under ComBat normalization setting. The red dots and associated values on the figure are the mean AUC scores for each method, while the black dots represent the outliers. Same method under different settings are represented in the same color of boxplots. All the experiments were repeated 30 times for each test dataset, and the results presented in the figure were based on the average AUC scores of the total 180 replications for the six test datasets.

**S2 Fig. Realistic applications of merging and integration methods on multiple tuberculosis genomic datasets**. The results by different methods are grouped into six groups. “Single learner”: Each of the five training datasets were trained independently with RF classifier and predicted on the test dataset, then the average AUC score was taken among the five predictions. “Merged”: Merging method with pooling all five training datasets into one train data. The “Single learner” and “Merged” experiments were conducted under both naive and ComBat normalization settings. “Ensemble learning”: The five training predictors were integrated by ensemble weighted learning methods under naive setting. “Ensemble learning (normalized)”: The five training predictors were integrated by ensemble weighted learning methods under ComBat normalization setting. “Rank aggregation”: The five training predictors were integrated by rank aggregation methods under naive setting. “Rank aggregation (normalized)”: The five training predictors were integrated by rank aggregation methods under ComBat normalization setting. The red dots and associated values on the figure are the mean AUC scores for each method, while the black dots represent the outliers. Same method under different settings are represented in the same color of boxplots. All the experiments were repeated 30 times for each test dataset, and the results presented in the figure were based on the average AUC scores of the total 180 replications for the six test datasets.

